# Quantitative Real-Time PCR assays for species-specific detection and quantification of Baltic Sea spring bloom dinoflagellates

**DOI:** 10.1101/871020

**Authors:** A. M. Brink, A. Kremp, E. Gorokhova

## Abstract

In the Baltic Sea, the dinoflagellates *Apocalathium malmogiense, Biecheleria baltica,* and *Gymnodinium corollarium* are important contributors to the spring bloom. However, their relative contribution to the bloom community cannot be unambiguously determined by conventional light microscopy due to lack of resolution of distinctive morphological features of the three species. Here, we describe a molecular approach based on a quantitative real-time polymerase chain reaction (qPCR) primer and probe system, targeting the ITS1 and ITS2 regions of the rRNA gene for all three species and enabling their quantification. The specificity of the method was demonstrated using monocultures of *A. malmogiense*, *B. baltica*, *G. corollarium* as well as three other dinoflagellate species co-occurring in the Baltic Sea during spring and validated using field-collected phytoplankton samples.

## INTRODUCTION

Dinoflagellates are arguably the largest group of marine eukaryotic phytoplankton aside from diatoms, and one of the most important primary producers in the marine ecosystem. Cold-water dinoflagellates are an important component of the algal spring bloom in the Baltic Sea (Lignell et al. 1993; Tallberg and Heiskanen 1998). During the last four decades, their proportion has increased significantly in some regions, and dinoflagellates now frequently dominate the spring bloom in the central and northern basins of the Baltic Sea (Klais et al. 2011; Spilling et al. 2018). Recent morphological and molecular analyses revealed that in addition to the chain-forming arctic *Peridiniella catenata*, at least three other species are associated with the spring dinoflagellate blooms in the Baltic (Larsen et al. 1995; Kremp et al. 2005; Sundström et al. 2009). The re-described *Biecheleria baltica* Moestrup, Lindberg et Daugbjerg, formerly known as *Woloszynskia halophila* (*sensu* Kremp et al., 2005) is common in the Gulf of Finland, co-occurring with *Apocalathium malmogiense* (G. Sjöstedt) Craveiro, Daugbjerg, Moestrup & Calado, formerly known as *Scrippsiella hangoei* (*sensu* Larsen et al., 1995) (Craveiro et al. 2017). The third species, *Gymnodinium corollarium* Sundström, Kremp et Daugbjerg (Sundström et al. 2009), occurs throughout the Baltic Sea and is particularly abundant in the open Baltic proper (Sundström et al. 2009).

Due to similar morphology, these species cannot be unambiguously distinguished from one another by light microscopy with or without staining, particularly in samples preserved with Lugol iodine solution (Kremp et al. 2005; Sundström et al. 2009). Unlike *B. baltica* and *G. corollarium*, *A. malmogiense* is armored; therefore, staining with CalcoFluor White could be applied to visualize the thecal plate patterns (Fritz and Triemer 1985). However, the thecal plates in this species are delicate and stain poorly, thus hampering the identification (Larsen et al. 1995). Cell surface features specific for unarmored *B. baltica* or *G. corollarium* can only be observed using scanning electron microscopy (SEM), a method not applicable for quantitative enumeration of algae in field plankton samples.

The difficulties with species identification are hampering our ability to study population dynamics of the individual dinoflagellate species and, thus, understand the seasonal succession of these ecologically important Baltic phytoplankton species. To understand whether phenology and bloom magnitude are driven by life-history traits and environmental factors involved in the life cycle regulation in *A. malmogiense, B. baltica,* and *G. corollarium* (Kremp et al. 2005), we need to distinguish these species during the analysis. Furthermore, variations in the relative contribution of these species to the spring bloom may affect their grazers. For example, interspecific differences in allocation of fatty acids (Leblond et al. 2006) may lead to higher egg production in copepods fed *G. corollarium* (Vehmaa et al. 2012) with consequences for energy transfer efficiency in the food web. Finally, species-specific encystment strategies might influence biogeochemical processes in the sediment once the bloom has settled out from the water column (Spilling and Lindström 2008); to study all these ecological processes, we need reliable methods for population analysis of these cold-water dinoflagellates co-occurring in the Baltic Sea.

In recent years, various molecular assays have been developed for the detection of several phytoplankton species, improving the detection capacity and reducing sample processing time. Quantitative polymerase chain reaction (qPCR) assays targeting ribosomal RNA (rRNA) encoding gene (rRNA operon) have become a popular tool for species identification and quantification in environmental samples of dinoflagellates (Bowers et al. 2000; Galluzzi et al. 2004; Dyhrman et al. 2006), particularly toxin-producing species with high capacity to outbreaks (Toebe et al. 2013; Smith et al. 2014; Kon et al. 2015; Hernández-Becerril et al. 2018; Engesmo et al. 2018; Ruvindy et al. 2018). In addition to high taxonomic resolution, these assays also have a sensitivity surpassing that of the microscopy-based techniques. For example, an assay developed for *Gymnodinium catenatum* estimated cell densities in environmental samples being as low as 0.07 cells per PCR reaction or three cells per liter and detected the presence of cells below the limit of detection for light microscopy (Toebe et al. 2013; Smith et al. 2014). Several types of real-time PCR assays with differing levels of specificity have been developed, and positive reactions are detected either with a fluorescent reporter probe (e.g., hydrolysis probes, molecular beacons, locked-nucleic acid bases [LNA]) or a double-stranded DNA-binding dye (e.g., SYBR green). As a target region for distinguishing dinoflagellate species, Internal Transcribed Spacer (ITS1 and ITS2) of the rRNA operon has been recommended as a high-resolution marker (Stern et al. 2012).

The aim of this study was to develop TaqMan qPCR assays applicable for identification and enumeration of *A. malmogiense*, *B. baltica*, and *G. corollarium* in the Baltic Sea plankton, with particular focus on the applicability of these assays in regular monitoring surveys. To ensure the assay specificity, each primer-probe set was tested not only with monocultures of *A. malmogiense*, *B. baltica*, *G. corollarium,* but also with three other dinoflagellate species co-occurring in the Baltic Sea with the target species during spring. The assay development proceeded in a step-wise manner and was validated using field-collected samples (**Figure 1**).

**Figure 1.**
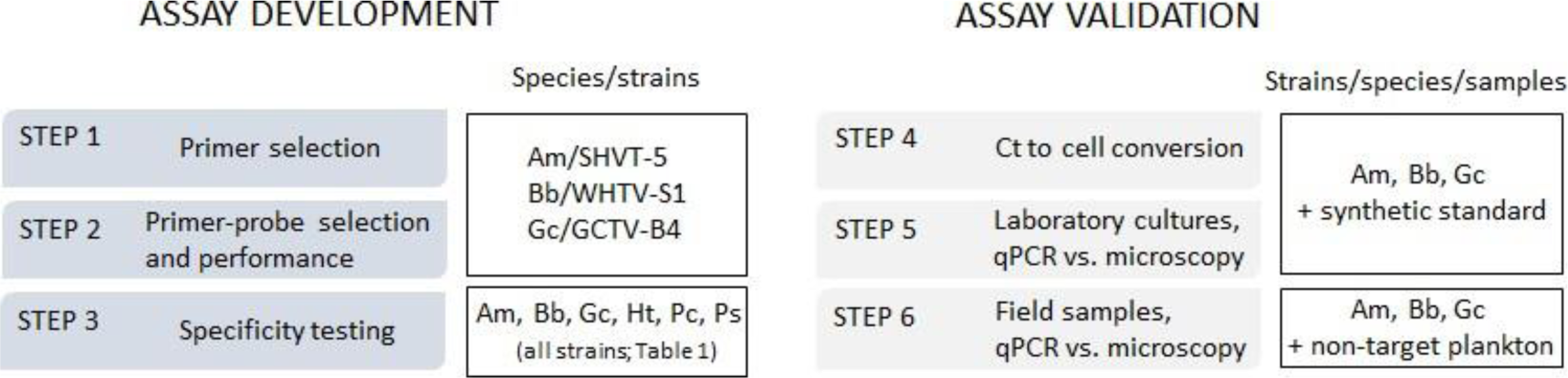
The outline of the qPCR assay development and validation steps and species/strains used for specific tasks. Species abbreviations: **Am** *-Apocalathium malmogiense,* **Bb** *-Biecheleria baltica,* and **Gc** *-Gymnodinium corollarium*.

## MATERIALS AND METHODS

### Test strains and environmental samples

#### Cultures

For the assay development, we used cultures of *A. malmogiense*, *B. baltica*, *G. corollarium,* and other dinoflagellate species that might be present in the plankton samples originated from the study area (**Table 1**, **Figure 1**). Except for *Protodinium simplex*, which was ordered from SCCAP (Denmark) and preserved with Lugol iodine solution upon arrival, all cultures used in this study (**Table 1**) were grown in temperature-regulated incubators at 50 μmol photons m^−2^ s^−1^ and a 14:10 h light:dark cycle. As a growth media, F/2-Si enrichment (Guillard and Ryther 1962) and salinity of 6.5 was used; throughout this paper, the salinity values are given in practical salinity units [PSU]. All cultures were grown at 4°C, except for *Heterocapsa triquetra*, which was maintained at 17°C. Subsamples of the cultures in the exponential growth phase were taken, preserved with Lugol iodine solution, and stored chilled in the dark until they were used for microscopy analysis and DNA extraction.

**Table 1.**
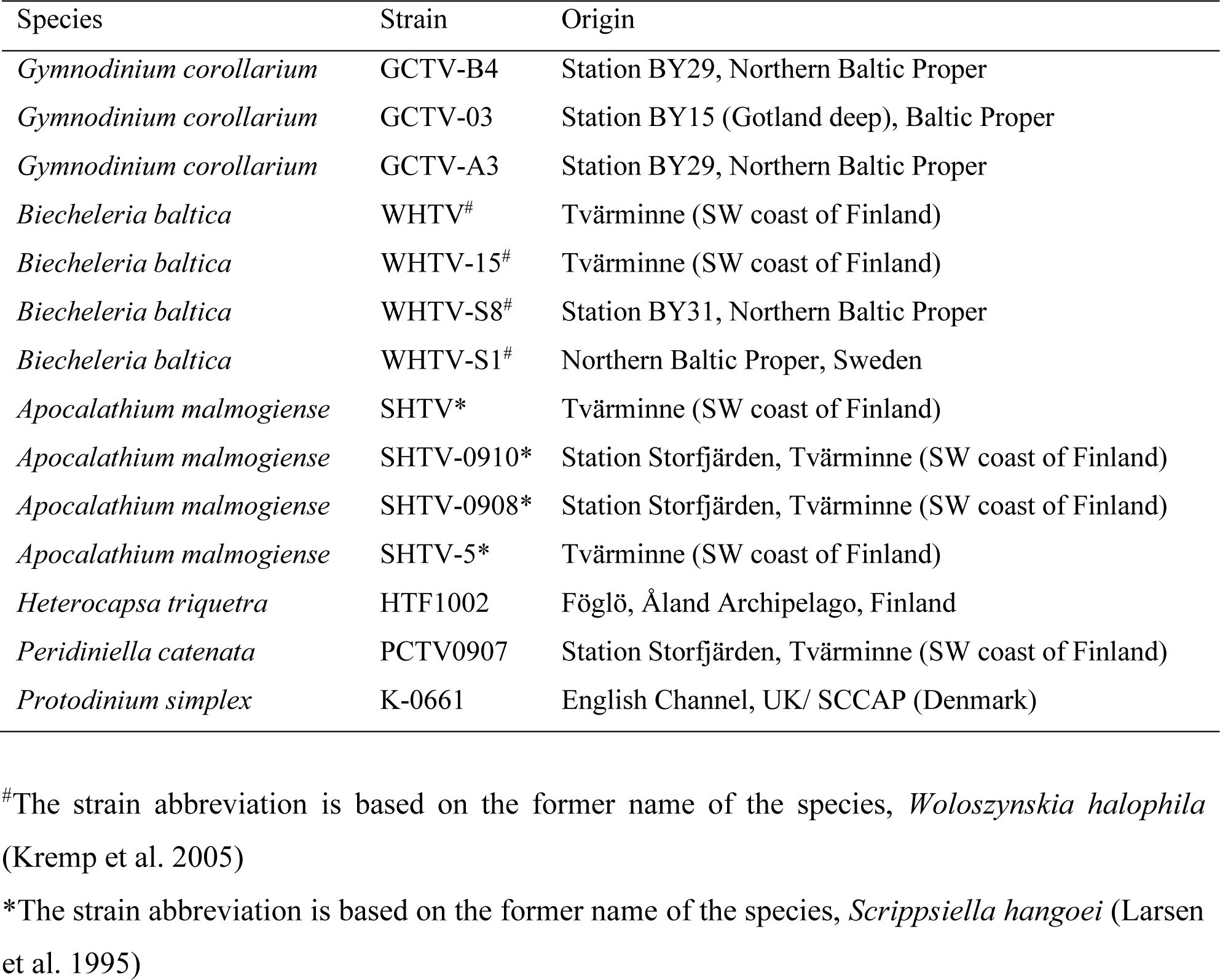
Cultures of dinoflagellate species and strains used in this study.

#### Field samples

In addition to the monocultures, field samples were collected in a coastal area (Himmerfjärden Bay, 58°.59’N, 17°.43’E; bottom depth 32 m) and used for the assay validation (**Figure 1**). Two time periods were selected for the sample collection to ensure (1) presence of abundant and diverse dinoflagellate communities (March 15–June 8, 2010), when all target species are present ubiquitously in the water column; and (2) absence of any of the target species in the plankton samples (September 28, 2010); by this time all of them should have undergone complete encystment and disappear from the pelagial (Kremp et al. 2009). All samples were collected during the daytime from the photic zone (0–14 m) using a plastic hose (inner diameter of 25 mm), and the entire sample volume was transferred to a bucket. After thorough mixing, two subsamples of the collected plankton were drawn from each sample and designated for: (1) microscopy analysis of the *Apocalathium/Biecheleria/Gymnodinium* complex; these samples (200 mL) were immediately preserved with 0.8 mL acetic Lugol iodine solution, and (2) qPCR assay; these samples (1 L) were preserved with 4 mL Lugol iodine solution. All samples were stored chilled in darkness until analyses.

### Microscopy analysis

Subsamples (25 mL) of the phytoplankton samples designated for microscopy-based identification and enumeration of dinoflagellates were settled for 20–40 h in Utermöhl chambers following the guidelines for phytoplankton analysis in the Baltic Sea (HELCOM 2017). Dinoflagellates were counted on the half-bottom or entire bottom of the sedimentation chamber under an inverted light microscope (Leitz DM-II) at 200× or 400× magnification. All counts were conducted in triplicate. Cell concentrations were calculated from obtained counts (cell mL^−1^) and used for the preparation of dilution series in qPCR assays, in the calculations of rRNA gene copy number per cell, and as an independent estimate of cell abundance.

### Design and verification of specific primers and probes

#### The qPCR approach

For each species, we designed primers and a hydrolysis probe (minor groove binder, MGB), which carries a fluorophore at the 5’-end and a quencher at the 3’-end. The hybridized probe is cleaved by the 5’ exonuclease activity of the Taq polymerase resulting in fluorescence increase. The species-specific primers and those probe sequences characterized by the highest number of mismatches to non-target sequences were tested *in silico* for their specificity by a BLAST sequence similarity search (http://www.ncbi.nlm.nih.gov/BLAST) against the GenBank nucleotide collection. Due to the additional specificity provided by the presence of the TaqMan probe, this approach was selected because it is less subject to false positives than the intercalating dye method (Kutyavin et al. 2000), which is crucial for mixed plankton samples.

#### Sequence information

The target for designing primers and probes was the rDNA ITS, regions ITS1 and ITS2. Sequencing of *B. baltica* (strain WHTV-C1) and *G. corollarium* (strain GCTV-B4) were performed in collaboration with the Department of Biomolecular Sciences, University of Urbino, Italy. The 5.8S rRNA gene and flanking ITS regions were amplified and sequenced using the BigDye Terminator Cycle Sequencing Kit (Applied Biosystems, Cheshire, UK) and an ABI PRISM 310 Genetic Analyzer instrument. The sequences have been deposited into the GenBank database (accession numbers MN525428 and MN525429 for *G. corollarium* GCTV-B4, and *B. baltica* WHTV-C1, respectively). For *B. baltica,* WHTV-C1 sequence and GenBank sequence DQ167868 were used to generate a consensus sequence. For *A. malmogiense*, a consensus sequence was created based on the GenBank sequences for the following strains: SHTV-2 (AY970655), SHTV-5 (AY970656), SHTV-6 (AY970657), SHTV-1 (AY499515), and 702 (EF205037).

#### qPCR primer and probe design

The primer-probe systems (**Table 2**) were first tested *in silico* for specificity. For each target species, alignments were made using the software BioEdit Sequence Alignment Editor version 7.0.9.0 (Hall 1999). Non-target species were included based on taxonomic relatedness to the target species, strong sequence match, and occurrence of the species in the Baltic Sea spring phytoplankton community. The alignments were searched manually to determine unique sequences within the ITS regions relevant for primer and probe design, which was then performed using Primer3 (Whitehead Institute and Howard Hughes Medical Institute, Maryland). A database similarity search was made using BLAST to ensure *in silico* specificity of primers and probes. Primers were synthesized by Sigma-Aldrich Sweden AB (Stockholm, Sweden) and probes by Applied Biosystems (Cheshire, UK). The probes were dual-labeled with the fluorophore 6FAM and the quencher MGBNFQ at the 5’ and 3’ ends, respectively.

**Table 2.**
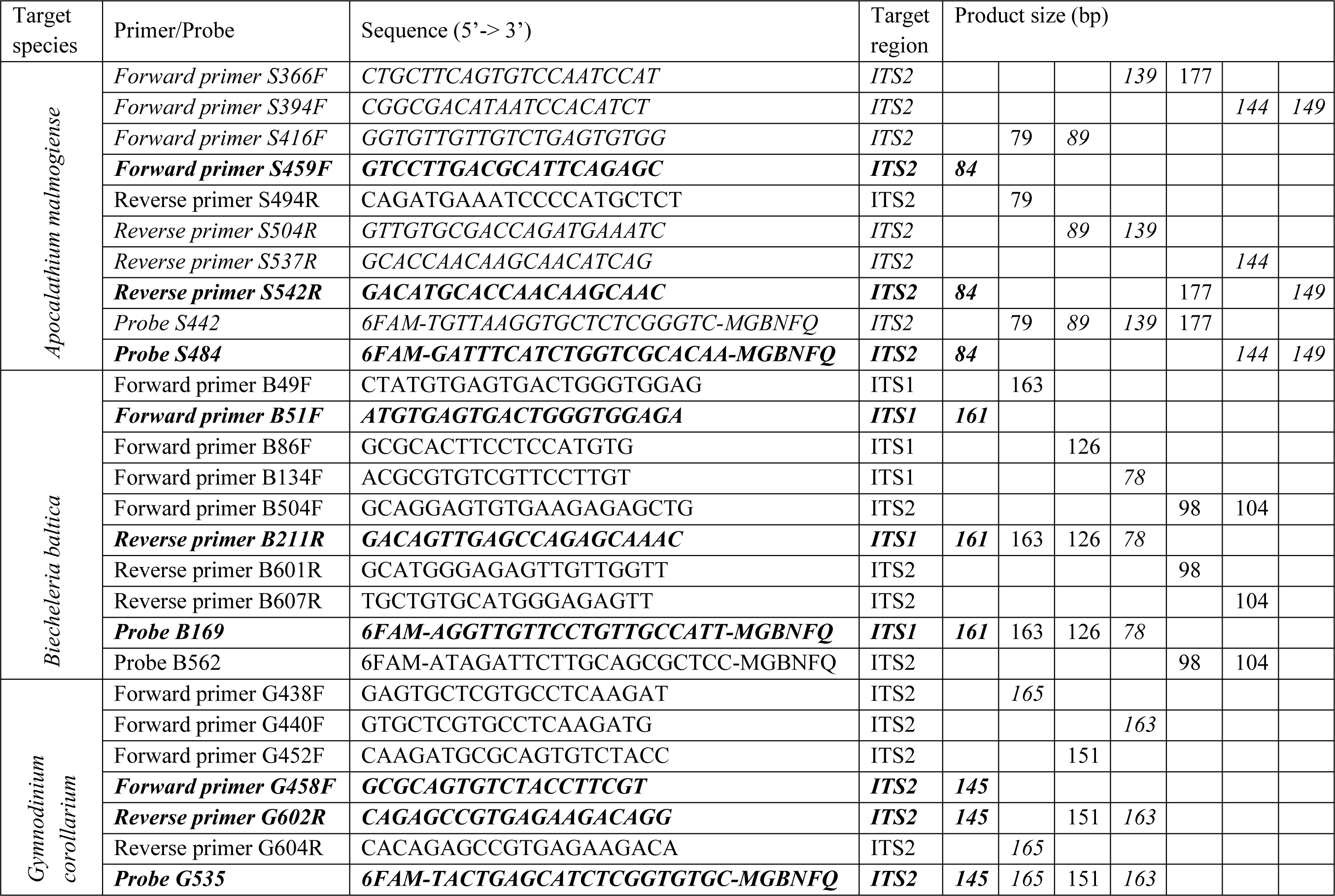
Primers and probes designed in this study, target regions and product size for the respective target species. Primers/probes chosen for qPCR testing are indicated in Italics. Primer/probe sets that were superior in the tests are indicated in bold.

### DNA extraction and quantification

#### DNA extraction

The DNA was obtained by Chelex extraction (Giraffa et al. 2000) using Lugol-preserved samples of dinoflagellates and wild plankton. For all extractions, the appropriate culture/sample volume was filtered onto a 5.0 µm Millipore TMTP Isopore polycarbonate membrane filter (Ø 25 mm) (Millipore, MA, USA) using a 15 mL Millipore filtration tower (Ø 16 mm) fitted to a vacuum filtration manifold. The filter was placed in a petri dish and cut diagonally with a scalpel into eight pieces which were all placed in a 2-mL tube. Nuclease-free water (200 µL; Qiagen, Hilden, Germany) was added, and samples were centrifuged at 10 000 rpm for 5 min and then frozen at −80°C. After thawing, each sample received approximately 100 mg of acid-washed glass beads (≤ 106 μm; Sigma-Aldrich, MO, USA) and was homogenized for 2 × 20 s in a FastPrep®-24 Instrument (MP Biomedicals, CA, USA). Then, 200 µL Chelex® 100, 10% w/v (Bio-Rad, CA, USA) were added followed by heating at 50°C for 30 min and then at 105°C for 8 min. The sample was centrifuged at 10 000 rpm for 5 min, the supernatant (100 µL) containing the DNA was transferred into a 1.5-mL tube and stored at 8°C overnight before processing.

#### DNA yield and purity assessment

Quantity and quality of the extracted DNA were evaluated by reading the whole absorption spectrum (220–750 nm) with a Nanophotometer^TM^ (Implen), and the DNA quality assessed using absorbance ratio at both 260/280 and 230/260 nm. Both ratios were good in most of the samples (1.8–2.0 and 2.0-2.2, respectively). In addition to the concentrations measured spectrophotometrically, the DNA concentration in each sample was also quantified fluorometrically by staining with Hoechst dye 33258 (Sigma-Aldrich) to avoid an overestimation of the DNA concentration measured by Nanophotometer (Bhat et al. 2010). The fluorometry-based DNA concentrations were applied to estimate template concentration in the TaqMan real-time PCR reaction.

### DNA standards and qPCR standard curves

#### Synthetic standards for qPCR

For all three species, gene-based standard curves were constructed to determine the efficiency of the assay, using a synthetic gene fragment approach (Vermeulen et al. 2009). A synthetic DNA oligonucleotide (Invitrogen Ltd.) comprising the target sequences (**Table 3**) and cloned into a plasmid was used as a standard (**Figure 2**). For each target species, the synthetic gene was assembled and cloned into pMA-T plasmid using *Sfil* and *Sfil* cloning sites. The plasmid DNA was purified from transformed bacteria (*E. coli*, K-12), and concentration was determined by UV spectroscopy. The final construct was verified by sequencing, and the congruence within the restriction site was 100%. For curves based on synthetic gene fragments, a 10-fold serial dilution ranging 75 to 75×10^5^ copies per reaction for all species of the plasmid construct containing the target sequence, were generated and used as a template. Each reaction was performed in duplicate.

**Figure 2.**
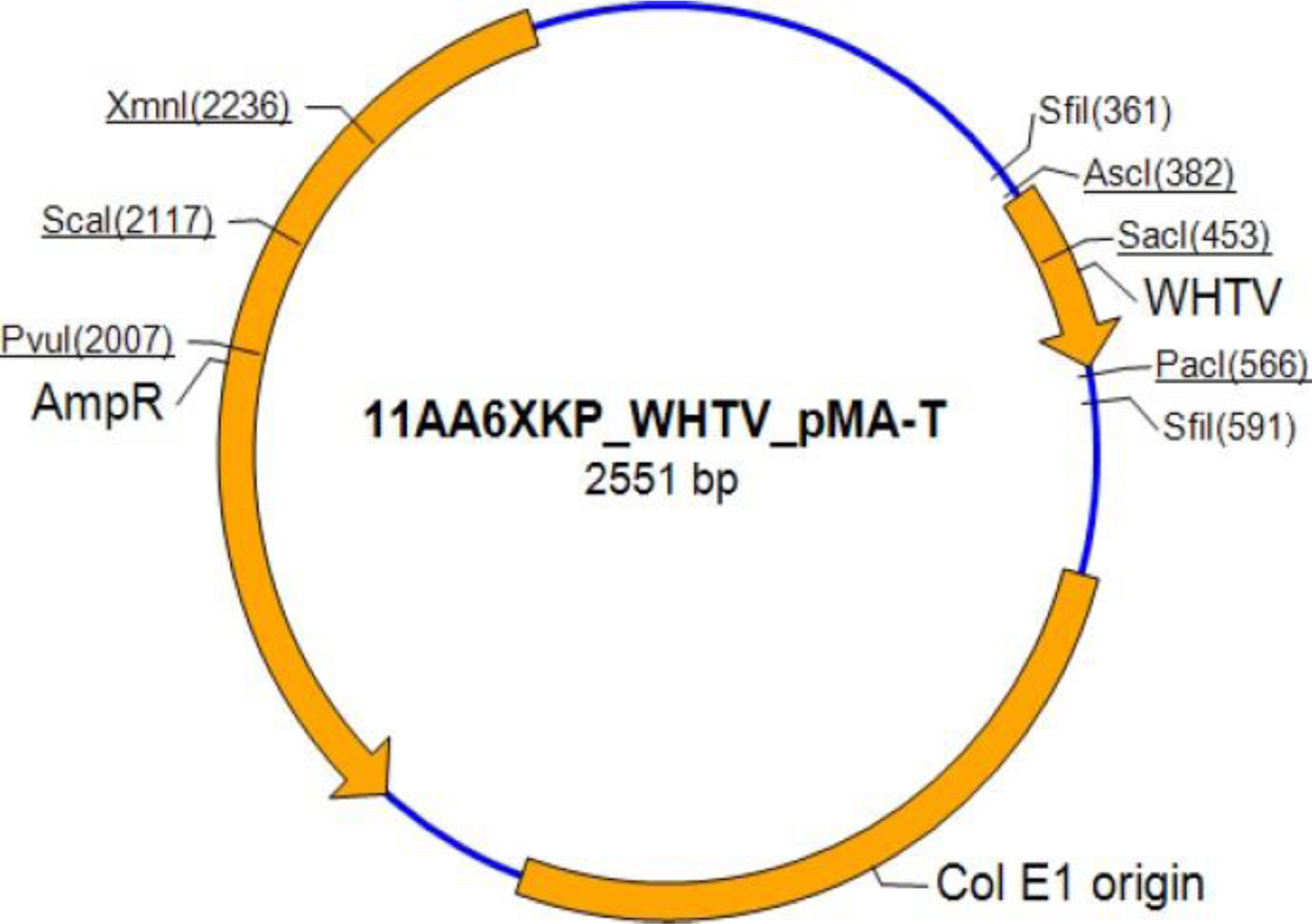
An example of pMA-T plasmid map for synthetic gene used for qPCR as a standard. Here, the construct corresponding to the target ITS1 fragment of *Biecheleria baltica,* strain WHTV, is shown. The plasmid maps for the other two target species were constructed in the same way using *Sfil* and *Sfil* cloning sites. The plasmid-inserted DNA was purified and lyophilized. When preparing the DNA for the standard dilutions, 5 µg of the plasmid DNA were dissolved in 50 µL distilled water.

**Table 3.**
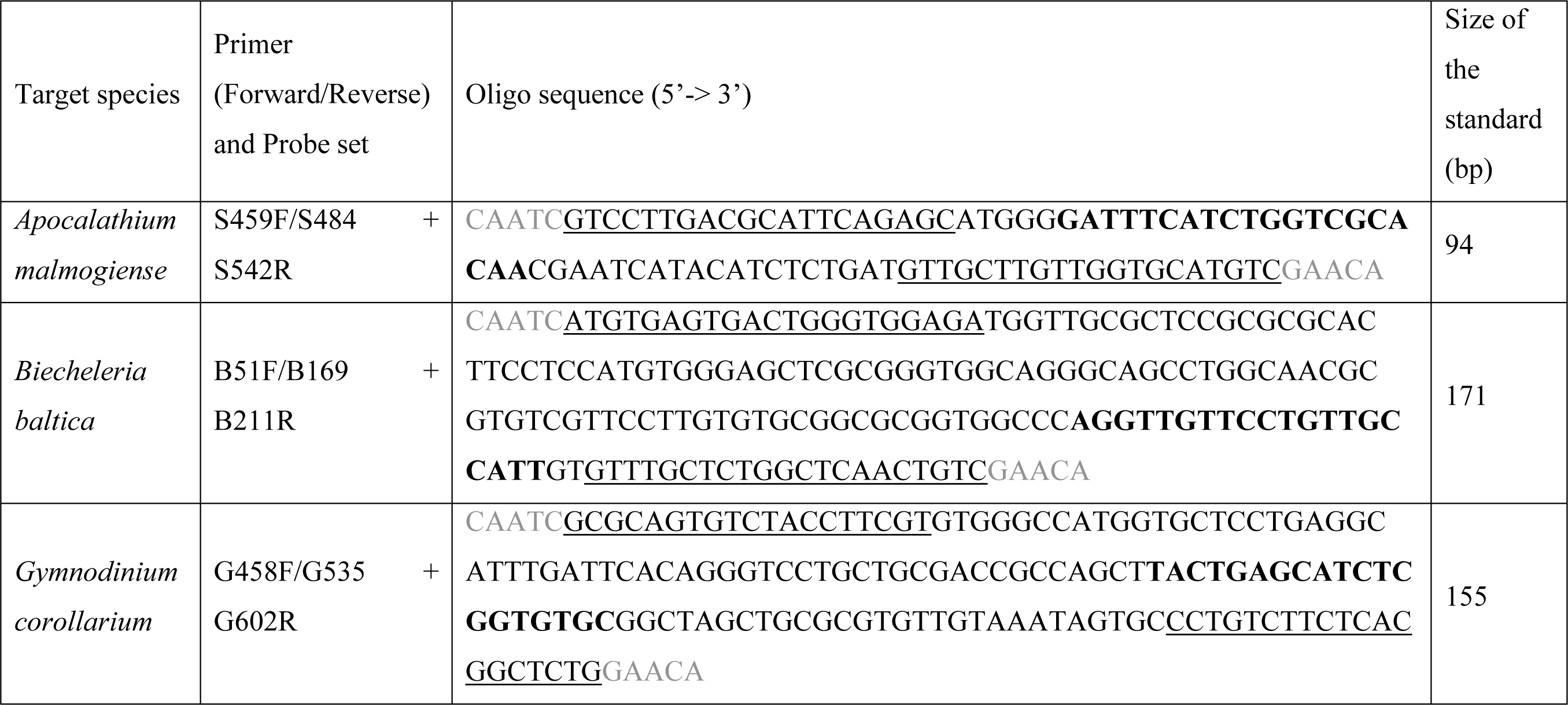
Synthetic oligonucleotides used as standards for *Apocalathium malmogiense*, *Biecheleria baltica,* and *Gymnodinium corollarium* in the qPCR assays. The oligo DNA includes sequence between the forward and the reverse primers (underlined) and the corresponding probe in-between (bold); CAATC and GAACA (grey letters) were added at the 5’ and 3’-end, respectively. The oligonucleotides were synthesized by Invitrogen (Life Technologies) and cloned into pMA-T plasmid (**Figure 2**).

#### Cell-based standard curves and determination of gene copy number per cell

One strain of each species, *A. malmogiense* (SHTV-5), *B. baltica* (WHTV-S1), and *G. corollarium* (GCTV-B4), were used to extract genomic DNA and construct cell-based standard curves. These extractions were serially diluted and used to generate target DNA representing a known cell number per reaction versus Ct data. The cell-based standard curves were constructed using 10-fold dilutions of the DNA extract using known cell concentrations ranging 1 to 8×10^3^ cells mL^−1^ for *A. malmogiense*, 1 to 7×10^3^ cells mL^−1^ for *B. baltica*, and 1 to 9×10^3^ cells mL^−1^ for *G. corollarium.* Each reaction was performed in triplicate.

#### Determination of gene copy number per cell

To determine the mean number of the rRNA gene copies per cell, the dilution series with a known cell count and gene-based Ct values were used as input for calculation. The slope of the linear regression between the rRNA gene copies and cell numbers in the reaction was used to determine the copy number per cell. This analysis was conducted for three strains per target species (**Table 1**).

### Assay performance

#### Primer performance

Candidate primers for *A. malmogiense*, *B. baltica,* and *G. corollarium* were tested using conventional PCR and DNA extracts from the monocultures of SHTV-5, WHTV-S1, and GCTV-B4, respectively. Each PCR reaction contained 2 µL of extracted DNA, 5 µL of Standard Taq Reaction Buffer (Final concentration 2.5×, 3.75 mM MgCl_2_, New England BioLabs, MA, USA), 1 µL of dNTPs (final concentration 0.5 mM, New England BioLabs), 0.5 µL of Taq polymerase (corresponding to 2.5 units, New England BioLabs), 2 µL of each primer (final concentrations 1 µM), and 7.5 µL of nuclease-free water (Qiagen, Hilden, Germany) resulting in a total reaction volume of 20 µL. For the non-template controls (NTC), nuclease-free water was used instead of DNA. Samples were amplified using MJ Research MiniCycler for 2 min at 94°C followed by 30 cycles of 15 s at 94°C, 15 s at 60°C and 45 s at 72°C with a final extension step for 7 min at 72°C. PCR products were mixed with 6× loading dye solution and ran on a 1.5 % agarose gel. GeneRuler 100 bp DNA Ladder (Thermo Scientific, MA, USA) was used as a marker. The gel was stained with ethidium bromide (0.2 µg/L) for 20 min, rinsed in distilled water for 7 min and visualized under UV light. All gels were examined to ensure that the PCR products had the expected size (bp) and no amplification occurred in NTC. Two main criteria for primer selection for qPCR were used: (1) efficient amplification as indicated by clear and sharp bands in the gels, and (2) the product size, with shorter amplicon size being favored as more suitable for qPCR (Penna and Galluzzi 2013).

#### Primer-probe performance

To ensure proper binding to the target species, functional primers with the corresponding probes were tested by qPCR using a StepOne Real-Time PCR instrument (Applied Biosystems). The following reagents were added for a 20 µL reaction mixture: 10 µL of TaqMan Gene Expression Master Mix (Applied Biosystems), 1.5 µL of each primer (0.75 µM), 0.5 µL of the probe (0.25 µM), 2 µL of the DNA extracted from a target species, and 4.5 µL of nuclease-free water. For the NTC, nuclease-free water was used instead of DNA. The thermal cycling conditions consisted of 2 min at 50°C and 10 min at 95°C followed by 40 cycles of 15 s at 95°C and 1 min at 60°C. Fluorescence data were collected at the end of each cycle, and the cycle threshold was determined automatically by the instrument software. Extracted DNA from the cultures SHVT-5, WHTV-S1 and GCTV-B4 was used in six-step tenfold serial dilutions (1×–10^5^×) and primer-probe sets were evaluated based on the qPCR efficiency and correlation coefficients (R^2^). All samples, standards, and NTC were analyzed in triplicates.

#### Reproducibility of the standard curve assays

The method’s reproducibility (inter-assay variation) was assayed by calculating the CV_Ct_ (coefficient of variation for cycle threshold) of gene- and cell-based standards in independent experiments run on different days using different sets of dilutions; five assays were used for gene-based standard curves and four assays were used for cell-based standard curves. The replicate standard curves were produced with the final set of primers and probes using a serial dilution series over the six-step tenfold serial dilutions (1×–10^5^×). The inter-run variation was calculated for C_t_ values as described by Rutledge and Côté (Rutledge and Côté 2003). For each primer-probe set the pooled slope and intercept was calculated and used for downstream applications of the gene-based standard curve.

#### Specificity testing

The specificity of each assay was confirmed by using DNA from the non-target species as a template. More specifically, cross-reactivity between primers and probe for *A. malmogiense* were tested with DNA extracted from the other two species, and the same was done for the assays developed for *B. baltica* and *G. corollarium*. The specificity of each primer-probe set was further tested by screening against non-target species *Heterocapsa triquetra*, *Peridiniella catenata* and *Protodinium simplex* (**Table 1**) that are phylogenetically close to the target species (Saldarriaga et al. 2001; Zhang et al. 2007). All primer-probe sets were also tested with DNA from a field sample collected in spring 2010—a time when all target species were expected to be present in the water column. The qPCR-products were sequenced (KIGene, Karolinska University Hospital, Stockholm, Sweden) to confirm the specificity of the assays. Furthermore, a field sample collected in the fall, not containing the target species, was also used in the qPCR assay. As positive controls, samples with the target species DNA were included in all assays.

### Statistical analysis

Statistics were performed with the GraphPad software, version 7 (GraphPad Software Inc.). The degree of linear relationship between data (R^2^ coefficient) was determined as the Pearson correlation coefficient and F-test was used to determine the overall significance of the regression. The regressions were compared with no assumption of the homogeneity of variances (Smithson 2012).

## RESULTS

### Optimization of primer-probe performance with conventional PCR and qPCR assays

#### Apocalathium malmogiense

Out of seven primer pairs (**Table 2**), one pair (S416F/S494R) produced an unspecific product in the NTC. All other pairs amplified the desired DNA region with no amplification in the NTC. However, the pair S366F/S542R was eliminated from further testing because of the relatively large amplicon, which was considered as suboptimal for qPCR. The remaining five pairs were combined with probes and tested in the qPCR assays. Among those, the best reaction efficiency (94.8) and R^2^ (∼1.00) were observed in the qPCR assay with primers S459F/S542R in combination with the probe S484.

#### Biecheleria baltica

Out of six primer pairs (**Table 2**), two pairs (B86F/B211R and B504F/B601R) produced unspecific products in the NTC and were eliminated from further testing. The remaining four pairs amplified the target region with no product in the NTC. Two of these, B51F/B211R and B134F/B211R were chosen for downstream qPCR testing, in combination with compatible probes (B169 in both cases). In the qPCR tests, the superior efficiency and R^2^ values, 93.3 and ∼1.00, respectively, were obtained for the primers B51F/B211R.

#### Gymnodinium corollarium

All four primer pairs (**Table 2**) amplified the desired target region with no amplification in the NTC. In the qPCR assay, the primer set G458F/G602R with G535 probe produced standard curves with the highest efficiency (96.9) and R^2^ = 0.99 (mean values for both parameters). In no case, a positive amplification was observed in the NTC of qPCR.

### Overall performance and reproducibility of the standard curve assays

For both gene- and cell-based assays, the standard curves were highly reproducible with high linear range (**Figure 3**), efficiency >90% (with a very few exceptions; **Table 4** and **Table 5**), and low inter-assay variation assayed by CV_Ct_ (**Table 6** and **Table 7**).

**Figure 3.**
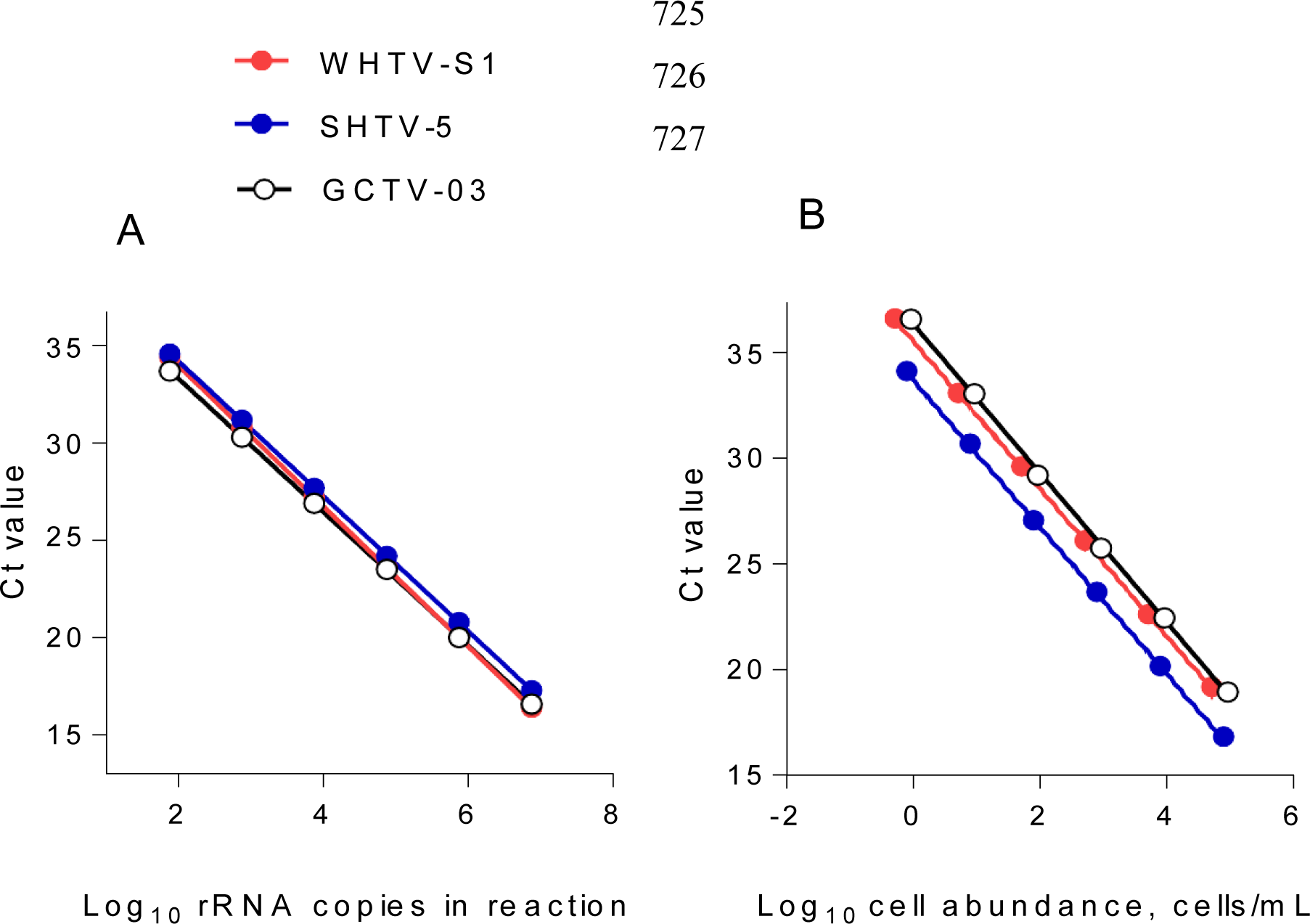
Standard curves of (A) Log_10_ starting rRNA gene copy number in the reaction vs. the threshold cycle number (Ct) for qPCR detection of DNA from synthetic controls in the gene-based assays (*n* = 6) and (B) cell-based assays for the target species/strains. All plots show mean values; the standard deviation bars are not visible (*n* = 5).

**Table 4.** Gene-based standard curves for *Apocalathium malmogiense*, *Biecheleria baltica*, and *Gymnodinium corollarium* generated on separate occasions (n = 5). Slope and intercept values are shown as mean ± SD. All assays included non-template controls that never produced a positive amplification. See also **Figure 3A**.

|  | Assay 1 | Assay 2 | Assay 3 | Assay 4 | Assay 5 |
| --- | --- | --- | --- | --- | --- |
| *Apocalathium malmogiense*, SHTV-5 |  |  |  |  |  |
| Slope | -3.55 ± 0.05 | -3.42 ± 0.08 | -3.46 ± 0.02 | -3.46 ± 0.03 | -3.43 ± 0.02 |
| Intercept | 41.57 ± 0.22 | 40.96 ± 0.37 | 41.25 ± 0.11 | 40.97 ± 0.18 | 40.84 ± 0.10 |
| Efficiency | 91.30 | 96.11 | 94.32 | 94.26 | 95.39 |
| R<sup>2</sup> | 0.998 | 0.995 | 1.000 | 1.000 | 1.000 |
| Pooled slope = -3.47 (p = 0.17). Pooled intercept = 41.14 (p = 0.07) |  |  |  |  |  |
| *Biecheleria baltica*, WHTV-S1 |  |  |  |  |  |
| Slope | -3.49 ± 0.02 | -3.51 ± 0.06 | -3.67 ± 0.07 | -3.54 ± .07 | -3.55 ± 0.12 |
| Intercept | 40.28 ± 0.14 | 40.90 ± 0.30 | 41.48 ± 0.35 | 41.04 ± 0.36 | 40.86 ± 0.60 |
| Efficiency | 93.31 | 92.58 | 87.29 | 91.55 | 91.23 |
| R<sup>2</sup> | 0.999 | 0.997 | 0.996 | 0.997 | 0.987 |
| Pooled slope = -3.60 (p = 0.37). Pooled intercept = 41.13 (p = 0,08) |  |  |  |  |  |
| *Gymnodinium corollarium*, GCTV-B4 |  |  |  |  |  |
| Slope | -3.41 ± 0.02 | -3.39 ± 0.07 | -3.47 ± 0.05 | -3.41 ± 0.02 | -3.41 ± 0.02 |
| Intercept | 40.05 ± 0.07 | 39.92 ± 0.28 | 40.29 ± 0.22 | 40.17 ± 0.12 | 40.23 ± 0.09 |
| Efficiency | 97.11 | 96.44 | 93.75 | 96.57 | 95.73 |
| R<sup>2</sup> | 1.000 | 0.998 | 0.999 | 0.999 | 1.000 |
| Pooled slope = -3.42 (p = 0.66). Pooled intercept = 40.14 (p = 0,08) |  |  |  |  |  |

**Table 5.** Cell-based standard curves for *Apocalathium malmogiense*, *Biecheleria baltica*, and *Gymnodinium corollarium* generated on separate occasions (*n* = 4). Slope and intercept values are shown as mean ± SD. All assays included non-template controls that never produced a positive amplification. See also **Figure 3B**.

|  | Assay 1 | Assay 2 | Assay 3 | Assay 4 |
| --- | --- | --- | --- | --- |
| *Apocalathium malmogiense*, SHTV-5 |  |  |  |  |
| Slope | $-3.47 \pm 0.03$ | $-3.49 \pm 0.03$ | $-3.47 \pm 0.01$ | $-3.45 \pm 0.02$ |
| Intercept | $33.50 \pm 0.09$ | $33.62 \pm 0.09$ | $33.79 \pm 0.04$ | $34.03 \pm 0.07$ |
| Efficiency | 94.21 | 93.43 | 94.21 | 94.96 |
| R<sup>2</sup> | 0.999 | 0.999 | 0.999 | 0.999 |
| *Biecheleria baltica*, WHTV-S1 |  |  |  |  |
| Slope | $-3.53 \pm 0.02$ | $-3.38 \pm 0.03$ | $-3.45 \pm 0.01$ | $-3.58 \pm 0.02$ |
| Intercept | $35.59 \pm 0.07$ | $35.66 \pm 0.09$ | $35.86 \pm 0.03$ | $35.23 \pm 0.07$ |
| Efficiency | 91.99 | 97.67 | 94.92 | 90.25 |
| R<sup>2</sup> | 0.999 | 0.999 | 1.000 | 0.999 |
| *Gymnodinium corollarium*, GCTV-B4 |  |  |  |  |
| Slope | $-3.44 \pm 0.03$ | $-3.65 \pm 0.07$ | $-3.48 \pm 0.07$ | $-3.53 \pm 0.02$ |
| Intercept | $35.87 \pm 0.10$ | $36.61 \pm 0.22$ | $36.31 \pm 0.21$ | $36.58 \pm 0.07$ |
| Efficiency | 95.30 | 87.92 | 93.84 | 91.85 |
| R<sup>2</sup> | 0.999 | 0.998 | 0.998 | 0.999 |

**Table 6.** Reproducibility of the qPCR assays based on standard curves with synthetic DNA dilutions (*n* = 5)

| Species/strain | Gene copies in the reaction | Mean Ct ± SD | CV<sub>Ct</sub> (%) |
| --- | --- | --- | --- |
| *Apocalathium malmogiense*<br>SHTV-5 | 7.5E+06 | 17.26 ± 0.17 | 1.02 |
| | 7.5E+05 | 20.79 ± 0.10 | 0.51 |
| | 7.5E+04 | 24.17 ± 0.15 | 0.65 |
| | 7.5E+03 | 27.67 ± 0.08 | 0.33 |
| | 7.5E+02 | 31.04 ± 0.36 | 1.16 |
| | 7.5E+01 | 34.57 ± 0.37 | 1.07 |
| *Biecheleria baltica*<br>WHTV-S1 | 7.5E+06 | 16.62 ± 0.27 | 1.60 |
| | 7.5E+05 | 20.03 ± 0.16 | 0.76 |
| | 7.5E+04 | 23.49 ± 0.22 | 0.94 |
| | 7.5E+03 | 26.92 ± 0.24 | 0.88 |
| | 7.5E+02 | 30.71 ± 0.30 | 0.98 |
| | 7.5E+01 | 34.73 ± 0.87 | 2.50 |
| *Gymnodinium corollarium*<br>GCTV-B4 | 7.5E+06 | 16.58 ± 0.19 | 1.12 |
| | 7.5E+05 | 20.01 ± 0.11 | 0.55 |
| | 7.5E+04 | 23.4 ± 0.14 | 0.62 |
| | 7.5E+03 | 26.96 ± 0.14 | 0.54 |
| | 7.5E+02 | 30.46 ± 0.28 | 0.93 |
| | 7.5E+01 | 33.60 ± 0.19 | 0.56 |

**Table 7.** Reproducibility of the qPCR assays based on standard curves with algal DNA dilutions (*n* = 4)

| Species/strain | Cell number in the reaction | Mean Ct ± SD | CV<sub>Ct</sub> (%) |
| --- | --- | --- | --- |
| *Apocalathium malmogiense*<br>SHTV-5 | 8.E+04 | 16.83 ± 0.21 | 1.26 |
| | 8.E+03 | 20.16 ± 0.27 | 1.37 |
| | 8.E+02 | 23.68 ± 0.36 | 1.53 |
| | 8.E+01 | 27.07 ± 0.29 | 1.10 |
| | 8.E+00 | 30.68 ± 0.28 | 0.93 |
| | 8.E-01 | 34.12 ± 0.12 | 0.37 |
| *Biecheleria baltica*<br>WHTV-S1 | 5.E+04 | 19.20 ± 0.62 | 3.26 |
| | 5.E+03 | 22.62 ± 0.48 | 2.16 |
| | 5.E+02 | 26.12 ± 0.55 | 2.11 |
| | 5.E+01 | 29.61 ± 0.37 | 1.28 |
| | 5.E+00 | 33.08 ± 0.41 | 1.27 |
| | 5.E-01 | 36.61 ± 0.21 | 0.58 |
| *Gymnodinium corollarium*<br>GCTV-B4 | 9.E+04 | 18.94 ± 0.17 | 0.91 |
| | 9.E+03 | 22.44 ± 0.37 | 1.67 |
| | 9.E+02 | 25.73 ± 0.18 | 0.71 |
| | 9.E+01 | 29.20 ± 0.29 | 1.02 |
| | 9.E+00 | 33.05 ± 0.25 | 0.78 |
| | 9.E-01 | 36.57 ± 0.42 | 1.16 |

#### Gene-based standard curves

The gene-based standard curves covered linear detection over six orders of magnitude, with a mean qPCR efficiency of 96%, 94% and 91% for *G. corollarium*, *A. malmogiense,* and *B. baltica*, respectively (**Table 4** and **Figure 3A**). The detection limit tested was 75 gene copy numbers in all assays. The Ct for the lowest gene copy number tested varied from 33.6 in GCTV to 34.7 in WHTV, so it is not likely that the sensitivity lower than 75 gene copy numbers can be achieved. The inter-assay variation was low, with CV_Ct_ mean values of the standard curves being 1.3%, 0.8% and 0.7% for *B. baltica*, *A. malmogiense,* and *G. corollarium,* respectively (**Table 6** and **Figure 4A**). The CV_Ct_ was not related to the copy number in the reaction in all assays. The pooled slope and intercept, calculated for the respective target species standard curve, are provided in **table 4**.

**Figure 4.**
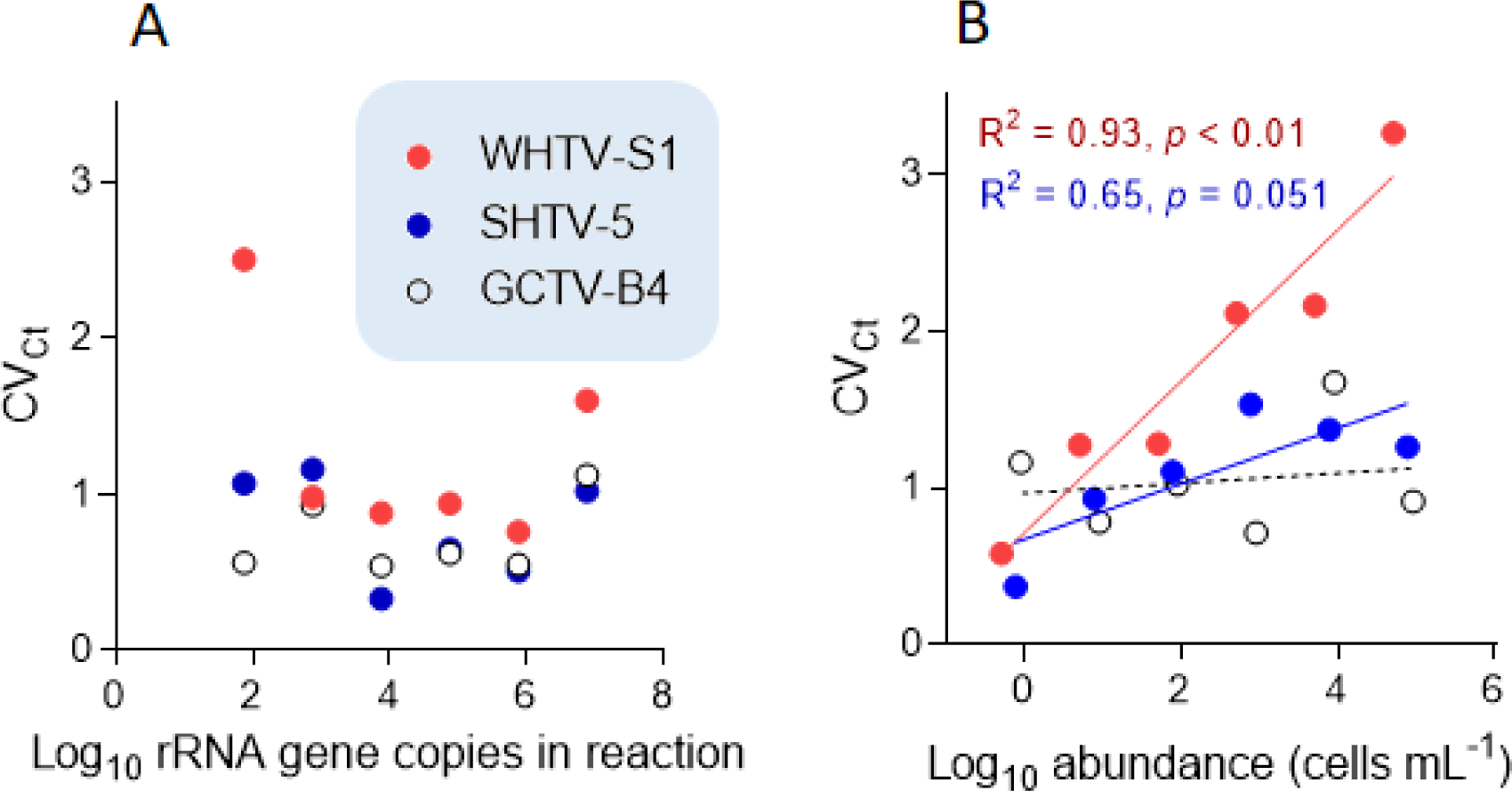
Inter-assay variation assessment for the target species and strains: *Apocalathium malmogiense* (SHTV-5), *Biecheleria baltica* (WHTV-S1), and *Gymnodinium corollarium* (GCTV-B4). Relationships between the CV_Ct_ values and (A) rRNA gene copy number in the reaction (*n* = 5) and (B) cell abundance in the cell-based qPCR assays (*n* = 5). Significant regressions for cell-based assays with WHTV-S1 and SHTV-5 are shown in the coding color. See **Table 4** and **Table 5** for the primary data for regression analysis.

#### Cell-based standard curves

The cell-based standard curves covered linear detection over six orders of magnitude, with a mean qPCR efficiency of 94% for *A. malmogiense* and *B. baltica* and 92% for *G. corollarium* (**Table 5** and **Figure 3B**). For all three species, the detection limit tested was less than one cell in the reaction. The CV_Ct_ mean values of the standard curves for the target species were 1.7%, 1.1% and 1.0% for *B. baltica*, *A. malmogiense* and *G. corollarium,* respectively, and remained at ≤1.0% for all species in the low cell abundance range (1 to 10^2^ cells mL^−1^). In *B. baltica*, the CV_Ct_ was significantly positively related to the cell abundance, whereas the relationship for *A. malmogiense* was nearly significant (**Figure 4**). In *G. corollarium*, the CV_Ct_ was the lowest among the three species (**Table 7**) and unrelated to the cell abundance (**Figure 4**).

### Evaluation of primer-probe specificity

No cross-reactivity for any of the laboratory strains of *A. malmogiense*, *B. baltica* and *G. corollarium* was observed (**Table 8**). Similarly, no amplification was observed for DNA extracted from the cultures of *Heterocapsa triquetra, Peridiniella catenata,* and *Protodinium simplex*, except for a few cases where non-specific product was observed after more than 35 cycles (the dynamic range of the assays; **Figure 3A**). Sequencing of the qPCR-products from the spring bloom field sample DNA confirmed the presence of all three target species and the specificity of the assays. Moreover, when the environmental DNA extracted from the field sample collected in September 2010 was used as a template, no specific amplification was observed with any of the primer-probe sets selected for the species-specific qPCR assays for *A. malmogiense*, *B. baltica* and *G. corollarium,* which was consistent with the absence of the species from the phytoplankton community at the time. However, for the *B. baltica* and *G. corollarium* primer-probe sets, some unspecific amplification occurred at Ct > 34, corresponding to the end of the standard curve and outside of the standard curve, respectively.

**Table 8.**
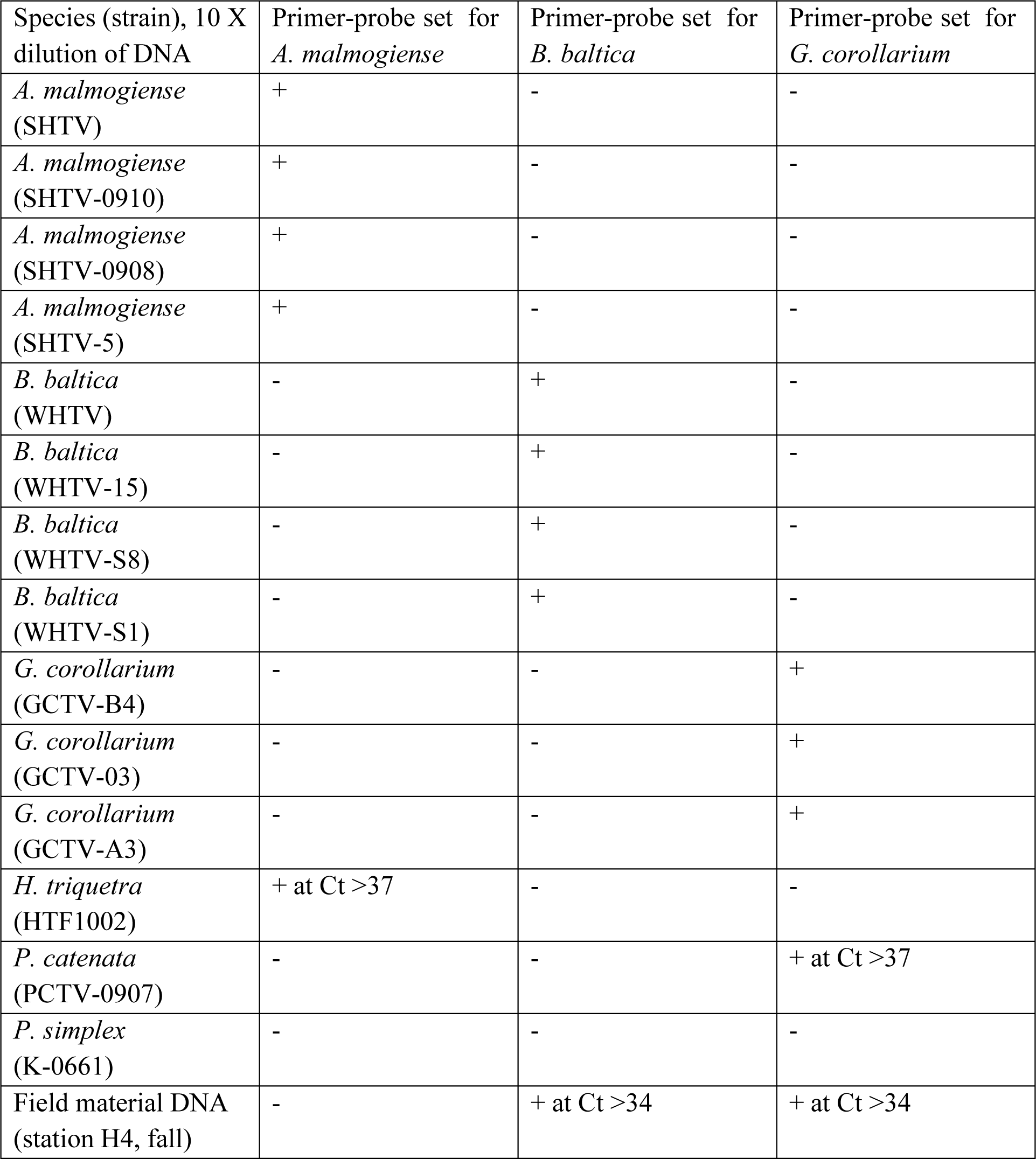
Cross-reactivity of the qPCR primer-probe sets for ecologically and phylogenetically relevant dinoflagellates included in the battery of test species/strains; see Table 1 for details. For non-specific amplifications, the Ct values are provided.

### Correspondence between microscopy- and qPCR-based estimates of cell abundance

There was very strong (R^2^ > 0.98 in all cases) correlation between the cell abundance estimates based on qPCR and the Utermöhl counts (**Table 9**). However, the qPCR assays consistently overestimated the number of cells in the culture samples used in the laboratory testing (**Figure 6**). Moreover, neither the slopes or the intercepts of the regression lines for the test strains were significantly different (*slopes*: F_2,12_ = 2.597, *p* > 0.12; the pooled slope equals 5.12; *intercepts*: F_2,14_ = 0.059, *p* > 0.94; the pooled intercept equals −217).

**Table 9.** Regressions for cell abundance estimates derived from the microscopy counts and qPCR assays for *Apocalathium malmogiense*, *Biecheleria baltica*, and *Gymnodinium corollarium.* Slope and intercept values are shown as mean ± SD; *n* = 3. See also **Figure 6.**

| Regression parameters | *A. malmogiense* | *B. baltica*, | *G. corollarium* | Pooled regression |
| --- | --- | --- | --- | --- |
| Slope | 5.66 ± 0.47 | 4.47 ± 0.32 | 5.13 ± 0.22 | 5.12 |
| Intercept | -1230 ± 1721 | 616.0 ± 1007 | -131 ± 900 | -217 |
| R<sup>2</sup> | 0.9728 | 0.9798 | 0.9927 | 0.999 |
| *p* value | 0.0003 | 0.0002 | < 0.0001 | < 0.0001 |
| Comparison of slopes and intercepts<br>(F- test) for all species | *Slopes:* F<sub>2, 12</sub> = 2.596; $p > 0.12$;<br>*Intercepts:* F<sub>2, 14</sub> = 0.058; $p > 0.9$ | | | |

### Screening environmental samples for target species

To evaluate the adequacy of the developed qPCR assays for environmental screening, they were applied to environmental community DNA extracts from plankton collected in a coastal area of the Northern Baltic Proper during spring bloom 2010 (**Figure 1**). For consistency, the pooled slope and intercept for the plasmid standard curve for each species (**Table 4**) were used when estimating cell abundances in the field samples. To convert the Ct values to the cell abundance estimates, we applied the average rRNA gene copy number observed for each species (**Figure 5**). Variable cell abundance was observed for the dinoflagellates comprising the *Apocalathium/Biecheleria/Gymnodinium* species complex, with clear peaks for *G. corollarium* and *B. baltica* (**Figure 7**). The most common species was *G. corollarum* detected at high abundance on all sampling occasions, whereas *A. malmogiense* was detected only twice and at low abundances. At moderate numbers, *B. baltica* was detected on all sampling occasions. The regression for the qPCR vs. microscopy was highly significant, although the qPCR-based estimates in most cases exceeded those based on the light microscopy (2.4-fold on average; **Figure 8**). Within the observed range of the dinoflagellate abundance (4 to 4×10^5^ cells L^−1^), there was a significant positive relationship for the fold-difference (between the qPCR and microscopy counts) and the total dinoflagellate abundance when the extreme value of 5.4-fold difference observed on the 15^th^ of March was excluded from the regression.

**Figure 5.**
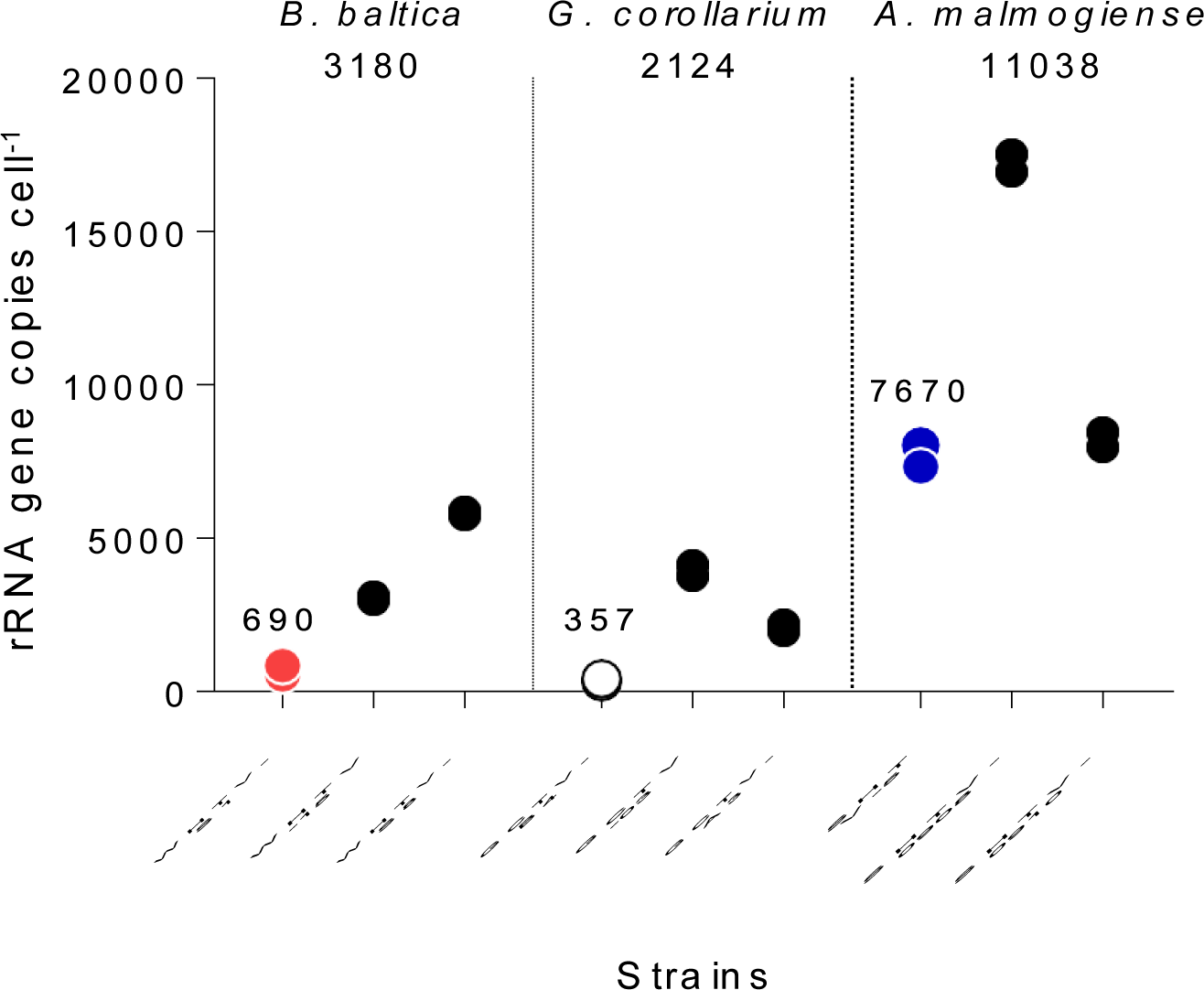
Variation in rRNA gene copy number per cell in the species and strains tested (**Table 1**). For each strain, two replicate samples were tested with three technical replicates. Mean value for all strains within a species is indicated below the species name and mean value for the strains used for the assay development (WHTV-S1, SHTV-5, and GCTV-B4) is indicated above the data points for these strains.

**Figure 6.**
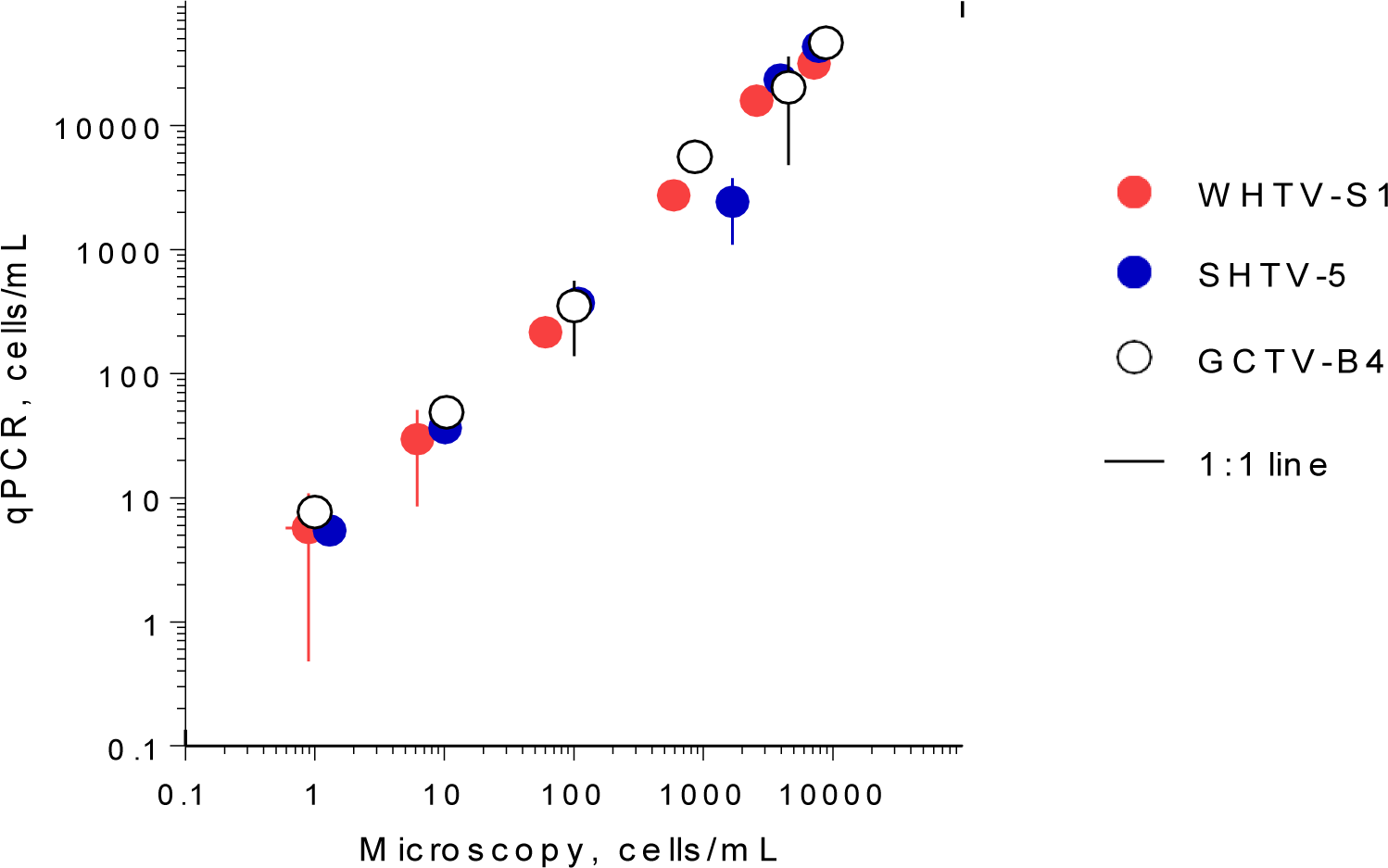
Correspondence for cell abundance estimates between microscopy- and qPCR based assays obtained for the test strains of the laboratory cultures of *Apocalathium malmogiense* (SHTV-5), *Biecheleria baltica* (WHTV-S1) and *Gymnodinium corollarium* (GCTV-B4). Horizontal and vertical error bars indicate 95% confidence interval based on three replicate measurements; note that in most cases, this variability is not visible. See Figure 5 for the strain-specific values of the rRNA gene copy number used in the calculations of cell abundance in the qPCR-based estimates and **Table 9** for statistical details on the linear regression for each species.

**Figure 7.**
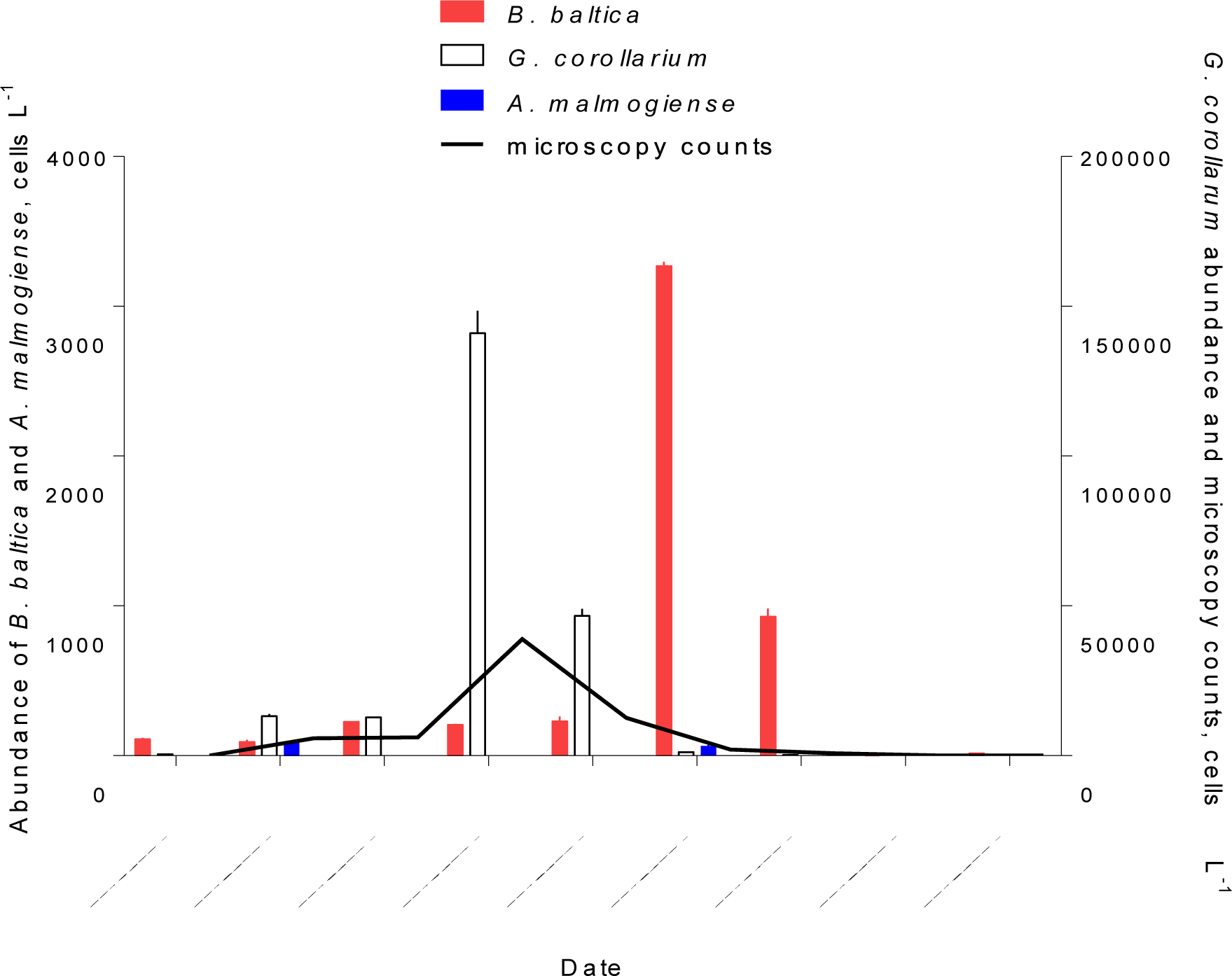
Application of the developed qPCR assays to field screening for the target species in the plankton samples collected in the Himmerfjärden Bay (H4) during the spring bloom period in 2010. For each species, a sample collected on each occasion was analyzed in triplicate using a respective synthetic standard and applying an average gene copy number determined for this species (Figure 5). The same samples were analyzed by light microscopy and the data for the entire species complex.

## DISCUSSION

In order to understand the dynamics of ecologically important algal blooms and to mitigate their impact on environmental and human health, it is important to optimize their monitoring in terms of frequency, sensitivity, and rapidity. Towards this goal, we need reliable and fast molecular methods with high sample throughput and low quantification thresholds. The identification of Baltic dinoflagellates in natural samples is currently based on light microscopy, but because of the low optical resolution that cannot visualize morphological differences among *Apocalathium malmogiense, Biecheleria baltica, and Gymnodinium corollarium*, species-level distinction is not possible. Moreover, even for experts, the identification is difficult, labor-intensive, and prone to errors, especially when several small-sized species co-occur in natural samples. To confirm the identification based on light microscopy, sophisticated equipment, such as SEM can be used, but it is not suitable for routine analysis of monitoring samples.

In the last decades, various molecular assays (e.g., real-time PCR, fluorescent hybridization assay, high resolution melting [HRM], sandwich hybridization assay, etc.) were shown to be instrumental for identification and enumeration of dinoflagellates. Earlier, a molecular identification method based on fluorescence in situ hybridization (FISH) assay has been developed for *B. baltica* (Sundström et al. 2010). The method was successfully applied to study the seasonal succession of this dinoflagellate at the SW coast of Finland. However, the labeling efficiency was affected by physiological changes in the algae during the bloom due to nutrient limitation, growth phase, and life cycle transitions; therefore, the quantification was prone to uncertainties (Sundström et al. 2010). Given the high specificity of the assays developed for many dinoflagellate species, a way forward for identification and quantification of the *Apocalathium/Biecheleria/Gymnodinium* complex would be applying the taxa-specific qPCR assays. Moreover, it is desirable that such assays would be applicable for regular plankton samples collected during monitoring and other surveys and preserved according to the existing monitoring practices (HELCOM 2017).

Here, we developed qPCR assays that can be used to rapidly detect co-occurring and morphologically similar dinoflagellates in plankton samples collected in the Baltic Sea. Species-specific qPCR primers and probes with high specificity and sensitivity were developed for three species, *A. malmogiense*, *B. baltica*, and *G. corollarium*. We also established that these primer-probe sets were effective in measuring the abundance of all three species in a coastal area of the Northern Baltic Proper. The cross-reactivity of the primers designed in this study showed high specificity for each target species while not amplifying when tested against other dinoflagellates reported from the study area. The species tested for cross-reactivity were chosen because they represented species that are genetically most similar to each target species for the ITS region; therefore, it is likely that the most likely candidates for false identification were included in the testing. Notably, the forward primer for *Apocalathium malmogiense*, S459F, also matches the ITS sequence of *A. aciculiferum* (Lemmermann) Craveiro, Daugbjerg, Moestrup & Calado (Craveiro et al. 2017). The reason is that *A. malmogiense* and *A. aciculiferum* have identical rDNA sequences and are believed to have diverged very recently (Logares et al. 2007). Therefore, it is likely that a positive amplification for *A. aciculiferum* can be obtained using the primer-probe system developed for *A. malmogiense*, but we have not tested it. However, besides having different morphological features compared to *A. malmogiense*, *A. aciculiferum* is a freshwater species inhabiting lakes, and it has not been reported in the marine environments, including the Baltic Sea. Therefore, we believe that for application in the Baltic, the selected system would be specific for *A. malmogiense*.

The mean amplification efficiencies for the developed gene- and cell-based assays targeting *G. corollarium*, *A. malmogiense* and *B. baltica* were >90%, with a 6-log linear dynamic range of quantification (**Table 4** and **Table 5**). The assays were sensitive as indicated by the fact that less than a single cell or 75 gene copies per reaction were sufficient to obtain a positive amplification. The quantification limit, below a single cell for all species (**Table 6** and **Table 7**), reflects the high rRNA gene copy number per cell. Indeed, as in other dinoflagellates, the rDNA operon is tandemly repeated up to thousands of copies (Saito et al. 2002; Galluzzi et al. 2004), which is supported by the observed variability in the rRNA gene copy numbers within and across the species (**Figure *5***). It is also possible that within-strain variability is also substantial depending on the life cycle and growth conditions as reported for several *Alexandrium* species in the Mediterranean Sea (Galluzzi et al. 2010).

Absolute quantification is usually determined by comparison to standard curves from target DNA using cell-based assays. However, the use of cell-based assays for quantification is complicated by the fact that dinoflagellates contain multiple copies of the rRNA gene (Guo et al. 2016). As we had no prior knowledge on the copy number variability between the target species and between different isolates of the same species, we used the synthetic gene standard and the developed qPCR assays to determine the relative difference in rDNA copy numbers between our target species and test strains (**Figure 5**). These data showed that there are indeed differences between the copy numbers both between species and between isolates within a single species, with the highest copy number observed in *A. malmogiense* and the lowest in *G. corollarium*. Moreover, the copy numbers varied by 8-, 11- and 2-fold, between different isolates of *B. baltica, G. corollarium*, and *A. malmogiense*, respectively. The rRNA cell copy number determined in this study were within the range of values reported for other dinoflagellates (Galluzzi et al. 2004, 2010; Penna and Galluzzi 2013).

Although we did not compare all dinoflagellate species listed in **Table 1**, the variability observed for the tested species and strains suggest that including more cultures in the cell copy number comparisons would increase the span of the observed values. Clearly, the use of cell-based standard curve would require more understanding of the ecological and evolutionary drivers of the rDNA operon variability for the species in question. To address these challenges, the synthetic gene approach is very instrumental and can be used for screening of field samples and determination of the absolute abundance of the target rRNA gene as well as the determination of cell copy number in specific strains and isolates. Given the observed cell copy numbers (**Figure 5**) and the dynamic range of the assays (**Table 6**. and **Table 7**), the sensitivity of the gene-based assays allows the analysis of less than a single cell in a qPCR reaction.

Accuracy of the absolute quantification approach relies on the quality of standard curve construction by controlling the precision and reproducibility (Rutledge and Côté 2003). A standard curve is generally considered to be of high quality if the correlation found between the log-copy numbers and the Ct values shows a coefficient superior to 0.99. Here, for all three species, the linearity was excellent (R^2^ > 0.99). However, the regression coefficient is a poor indicator of the precision or accuracy achieved (Rutledge and Côté 2003). In the comparison of Ct values among five independent runs of the cell-based assays, we found an average CV_Ct_ variation of <2% for all species (**Table 7** and **Figure 4B**), which appeared to indicate a good reproducibly between runs. Moreover, in the low range of the target cells, the CV_Ct_ values were below 1% (Table 4), indicating that at the environmentally relevant concentrations of the dinoflagellates, the assays are highly reproducible. As expected, the inter-assay variation for the gene-based assays was even lower (CV_Ct_ <1%; **Table 6** and **Figure 4A**), emphasizing usefulness of the synthetic standard for internal quality control of the assay reproducibility (Vermeulen et al. 2009; Conte et al. 2018).

A correspondence was found between the results obtained with microscopy and those found with each of the qPCR assays applied to the single-species cultures (**Figure 6**). This high correlation was also evident when the abundance of the entire *Apocalathium/Biecheleria/Gymnodinium* complex was estimated during the bloom event in a coastal area of the Northern Baltic Proper (**Figure 7** and **Figure 8**). However, the qPCR assays consistently overestimated the cell abundance, both in the laboratory tests with monocultures (5-fold; **Figure 6**) and in the mixed field-sampled plankton (2-fold; **Figure 8**). A difference of this magnitude would lead to considerably different abundance estimates of the target species. Moreover, the discrepancy between the qPCR-based estimates and the microscopy counts increased significantly with increasing cell abundance, indicating that during the peak bloom, the uncertainty would be the highest. The mounting evidence of intra-strain variability in detectable rDNA copy numbers and growth rate could have severe implications for qPCR-based cell enumeration of dinoflagellates, but also other groups, such as diatoms (Guo et al. 2016), and requires further investigation.

**Figure 8.**
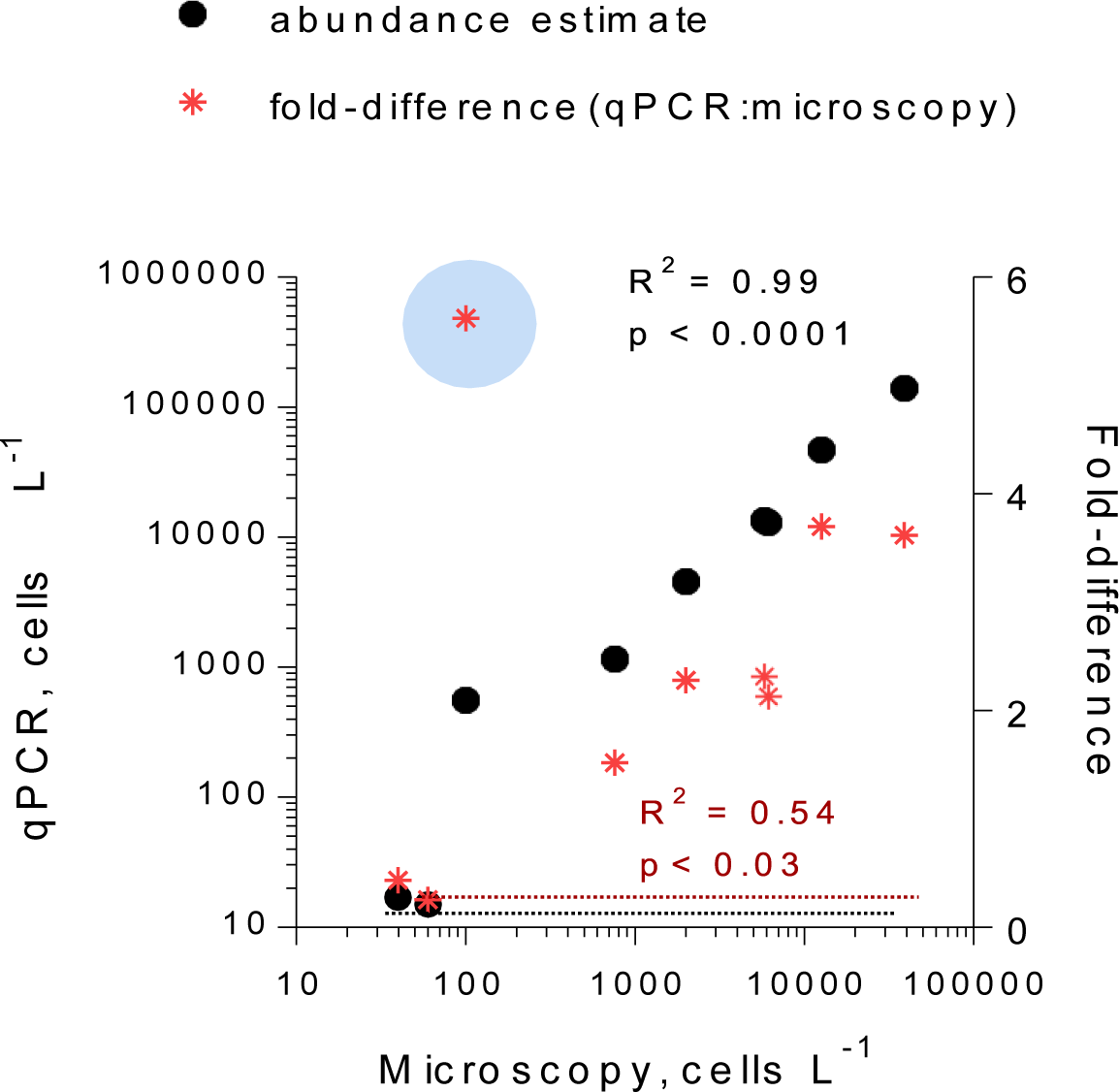
Correspondence for the total abundance (cells L^−1^) of the dinoflagellates belonging to *Apocalathium/Biecheleria/Gymnodinium* complex between estimates derived from the microscopy counts (no species-level identification is possible) and the qPCR assays targeting each species. The regression for the qPCR vs. microscopy was based on all data points, whereas the extreme value of 5.4 (indicated with a blue circle) was excluded from the regression for the fold-difference as a function of microscopy counts. See Figure 7 for the primary data.

In conclusion, this study demonstrates the usefulness of real-time PCR as a sensitive and rapid molecular technique for the detection and quantification of *A. malmogiense*, *B. baltica*, and *G. corollarium* from environmental samples. For each species, the inter-assay variation of the cell-based assays was low, with CV_Ct_ <1% in the ecologically relevant range of population abundance, which facilitates their applicability in the analysis of the monitoring or other samples that are collected and analyzed continuously. The assays developed are highly specific and sensitive in the unambiguous detection of all three species, and thus are valuable for routine plankton, biogeographic and phylogenetic investigations. Future studies should address ecological and phylogenetic aspects of rRNA copy number variability and improve its assessment in the monitoring and field studies of *A. malmogiense*, *B. baltica*, and *G. corollarium*.

## ACKNOWLEDGEMENTS

Laboratory facilities were provided by the Department of Ecology, Environment and Plant Sciences (Stockholm University). We thank Prof. Antonella Penna and her co-workers at the Department of Biomolecular Sciences (University of Urbino) for assisting with the sequencing work. This work was partly funded by the Walter and Andrée de Nottbeck Foundation and the Åland autonomy anniversary fund.

